# Epitranscriptome-wide profiling identifies RNA editing events regulated by ADAR1 that are associated with DNA repair mechanisms in human TK6 cells

**DOI:** 10.1101/2025.07.11.664482

**Authors:** Akito Yoshida, Yuqian Song, Hotaru Takaine, Sujin Song, Yu-Hsien Hwang-Fu, Zachary Johnson, Kiyoe Ura, Akira Sassa

## Abstract

Adenosine-to-inosine (A-to-I) editing is an endogenous RNA modification in eukaryotes, catalyzed by adenosine deaminases acting on RNA (ADARs). This modification modulates the gene expression by influencing splicing, RNA stability, and coding potential, depending on the site of editing. Although recent studies suggest a crosstalk between A-to-I editing and transcripts involved in DNA repair, the extent and functional significance of this interaction remain unclear. To investigate this, we applied the EpiPlex RNA assay—a method enabling epitranscriptome-wide detection of RNA modifications—in human lymphoblastoid TK6 cells. Across two biological replicates, we identified 869 transcripts bearing A-to-I–modifications. Gene Ontology analysis revealed significant enrichment in genome maintenance pathways, including “chromatin remodeling” and “DNA repair.” Notably, 27 transcripts encoding DNA repair proteins—such as *ATM, FANCA, BRCA1, POLH*, and *XPA*—contained A-to-I sites within introns or 3′ untranslated regions. To assess the isoform-specific contributions of ADAR enzymes—specifically ADAR1 p150 and p110—to RNA editing, we generated p150-deficient (p150 KO) and p150/p110-deficient (p150/p110 KO) TK6 cells. A-to-I editing peaks were reduced by ∼73.4% in p150 KO cells and nearly abolished (99.9%) in p150/p110 KO cells, indicating that most editing sites are p150-dependent, while a notable subset relies on p110. Importantly, a novel splice variant of *XPA* emerged in ADAR1-deficient cells, suggesting a role for RNA editing in alternative splicing regulation. Our epitranscriptomic analysis of A-to-I RNA editing underscores a multifaceted role for ADAR1-dependent editing in preserving genome integrity through posttranscriptional regulation of DNA repair genes, laying the groundwork for future studies into RNA-based mechanisms of genome maintenance.

## 1 Introduction

RNA modifications are post-transcriptional chemical alterations of RNA nucleobases that play crucial roles in gene regulation in mammals. Major RNA modifications such as *N*^6^-methyladenosine (m6A), 5-methylcytidine (5-mC), pseudouridine (Ψ), Adenosine-to-Inosine (A-to-I) editing, dynamically regulate RNA metabolism, including splicing, nuclear export, stability, translation, and degradation (1). Among these, A-to-I editing is a unidirectional RNA modification catalyzed by Adenosine Deaminase Acting on RNA (ADAR), which specifically convert adenosine-to-inosine within double-stranded RNA (dsRNA) regions (2). In humans, the ADAR family comprises three members: *ADAR1, ADAR2*, and the catalytically inactive *ADAR3*. While *ADAR2* is predominantly expressed in the central nervous system, *ADAR1* is broadly expressed across various tissues. ADAR1 exists in two major isoforms generated through alternative splicing: the constitutively expressed nuclear isoform *ADAR1* p110, and the interferon-inducible cytoplasmic isoform *ADAR1* p150, which contains a Z-DNA-binding domain (3, 4). Notably, ADAR1 p150 prevents inappropriate activation of innate immune signaling pathways by editing cytoplasmic dsRNAs (5-7). In addition to its enzymatic activity, ADAR1 also functions as an RNA-binding scaffold protein independent of A-to-I-editing. For instance, it directly interacts with Dicer to facilitate pre-miRNA processing and enhance miRNA loading into the RNA-induced silencing complex, highlighting its role in RNA interference (8). Furthermore, ADAR1 p110 has been shown to stabilize antiapoptotic mRNAs following UV irradiation by inhibiting Staufen1-mediated mRNA decay, thereby supporting cell survival (9).

Advancements in next-generation sequencing technologies have enabled transcriptome-wide identification of A-to-I editing sites (10, 11). The functional impact of A-to-I editing varies depending on the location of the modification within the mRNA (12). A significant proportion of editing occurs in the 3′ untranslated regions (3′UTRs) and introns, where dsRNA structures formed by Alu repeat elements serve as editing substrates (13-15). Intronic editing can influence alternative splicing by creating or abolishing splice sites, as observed in genes such as *NARF* and *CDDC15* (16, 17). Editing within 3′UTRs can disrupt miRNA binding, thereby modulating mRNA stability and translational efficiency. For example, ADAR1-mediated editing of *DHFR* and *METTL3* mRNAs has been linked to increased expression of these genes and tumor progression in breast cancer cells (18, 19). A-to-I editing can also occurs within coding regions, leading to amino acid substitutions that may alter protein function. ADAR2-mediated editing of the AMPA receptor subunit GluA2 exemplifies this, as it controls calcium permeability and suppresses excitotoxicity in neurons (20). Additionally, A-to-I editing of lncRNAs can affect RNA secondary structure and miRNA binding, thereby regulating gene expression. In the case of NEAT1, such modifications enhance RNA folding stability and prevent competition with miR-9-5p, ultimately modulating JAK-STAT signaling and interferon-responsive genes expression (21).

Recent studies have proposed that A-to-I editing plays a regulatory role in the DNA damage response and DNA repair pathways. For example, ADAR1-mediated editing within the exon of *NEIL1* mRNA, which encodes a DNA glycosylase, results in a K242R amino acid substitution. The edited NEIL1 protein exhibits diminished capacity to recognize and excise damaged bases (22, 23). A-to-I editing has also been reported in the 3′UTRs of *BRCA2, ATM*, and *POLH* mRNAs, where increased editing correlates with transcript expression levels (24). In the case of *BRCA2*, editing prevents miRNA binding, thereby stabilizing the mRNA and enhancing protein levels, contributing to cisplatin resistance (25). These findings highlight the need for systematic investigation into the interplay between A-to-I editing and genome maintenance, particularly through factors involved in DNA repair and the DNA damage responses. To address this, we aimed to comprehensively identify inosine-modified transcripts associated with DNA repair pathways.

To explore the potential crosstalk between A-to-I RNA editing and DNA repair factor transcripts, we focused on the human lymphoblastoid cell line TK6, which is extensively used in genome stability and genotoxicity research. TK6 cells retain functional p53 and exhibit a stable karyotype, making them a reliable model for studying the relationship between RNA editing and genome maintenance. We employed the EpiPlex RNA assay (26), a technique that enables epitranscriptome-wide mapping of both m^6^A and inosine modifications in cellular RNAs by enriching for modifications in fragmented RNA, to perform a comprehensive analysis of RNA modification dynamics. The resulting epitranscriptomic mapping revealed significant enrichment of A-to-I editing within DNA repair-related pathways. Leveraging this dataset, we further investigated the isoform-specific roles of ADAR1 by comparing inosine modifications in DNA repair transcripts under conditions deficient in either ADAR1 p110 or p150. Notably, the nucleotide excision repair factor XPA was found to undergo splicing regulation via intronic inosines. This study provides new insights into the regulatory role of RNA inosine modification in DNA repair processes.

## 2 Materials and Methods

### 2.1 Cell culture

The human lymphoblastoid The TK6-derived human lymphoblastoid cell line TSCE5 (referred to hereafter as TK6 for simplicity) were cultured in RPMI-1640 medium (Nacalai Tesque) supplemented with 200 μg/mL sodium pyruvate, 100 U/mL penicillin, and 100 μg/mL streptomycin, and 10% (v/v) heat-inactivated fetal bovine serum (Nichirei Biosciences, Inc.) (27). The cultures were maintained at 37°C in a 5% CO_2_ atmosphere with 100% humidity.

### 2.2 Generation of *ADAR1* deficient TK6 cell lines

To generate *ADAR1* p150 and p110 deficient cells, we designed single-guide RNA (sgRNA) targets for CRISPR/Cas9 genome editing in combination with gene targeting constructs. Primers used in the experiments are listed in Supplementary Table S1. CRISPR-target sequences are depicted in Figure 2B. sgRNAs were inserted into the *Bbs*I site of the pX330 vector. For disruption of *ADAR1* p150, the p150 targeting plasmid was constructed as follows; left and right arm fragments were PCR-amplified from TK6 genomic DNA. The amplified fragments were assembled via a seamless reaction (GeneArt® Seamless Cloning and Assembly Kit; Invitrogen) into the DT-ApA/PURO^R^ vector (kindly gifted by Dr. Hiroyuki Sasanuma), which is predigested with *Apa*I and *Afl*II. The vectors pX330-gRNA/p150 (6 μg) and the p150 target plasmid (2 μg) were transfected into TK6 cells by a NEPA21 electroporator (Nepa Gene Co. Ltd.) following the manufacturer’s instructions. After 48h incubation, cells were seeded into 96 microwell plates in the presence of puromycin (0.5µg/ml). Drug-resistant cell colonies were picked 10-14 days after transfection and subjected to genomic PCR for a targeted allele with puromycin-resistance cassette. For disruption of *ADAR1* p110, the p110 target plasmid was constructed as follows; left and right arm fragments were PCR-amplified from TK6 genomic DNA. The DNA fragment containing the neomycin resistance marker gene conjugated with the simian virus 40 polyadenylation signal (*NEO*^R^-polyA) was amplified from pUC-NSD2-*NEO*^R^ (28). The resulting left arm, right arm, and the *NEO*^R^-polyA fragments were assembled via a seamless reaction (GeneArt® Seamless Cloning and Assembly Kit; Invitrogen). The vectors pX330-gRNA/p110 (6 μg), and the p110 target plasmid (2 μg) were then transfected into cells by a NEPA21 electroporator (Nepa Gene Co. Ltd.). After 48h incubation, cells were seeded into 96 microwell plates in the presence of neomycin (1.0 mg/ml). Drug-resistant cell colonies were picked 10-14 days after transfection and subjected to genomic PCR for a targeted allele with neomycin-resistance cassette of p150/p110 KO construct.

### 2.3 RT-PCR

Total RNA was extracted from cells using the NucleoSpin RNA Plus kit (Macherey-Nagel). cDNAs were synthesized from the extracted RNA with ReverTra Ace (Toyobo Co., Ltd.). The RNA was then reverse transcribed using ReverTra Ace® (Toyobo, Co., Ltd.). Quantitative PCR was performed using Thunderbird® Next SYBR qPCR Mix (Toyobo, Co., Ltd.) and specific primers listed in Supplementary Table S1. The expression levels of the genes were normalized to the internal *GAPDH* expression levels. To amplify the XPA fragments spanning exons 2–6 and exons 5–6, PCR was performed using KOD-FX-Neo (Toyobo, Co., Ltd.) and cDNA derived from the cell lines with respective primers listed in Supplementary Table S1.

### 2.4 Western blotting

Total cell extracts were fractioned on 10% SDS-polyacrylamide gels and transferred onto PVDF membranes. Membranes were blocked with 3% skim milk before incubation with primary antibodies. To detect ADAR1 and GAPDH, membranes were incubated overnight at 4°C in Hikari A solution (Nacalai Tesque) or 3% skim milk with 1:1000 dilution of anti-ADAR1 monoclonal antibody (D7E2M, Cell Signaling) or 1:1000 dilution of anti-GAPDH monoclonal antibody (sc-32233, Santa Cruz), respectively. After washing with Tris-buffered saline containing 0.05% Tween 20, the membranes were incubated with a 1:4000 dilution of anti-mouse IgG conjugated to horseradish peroxidase (Cytiva) in Hikari B solution (Nacalai Tesque) or 3% skim milk. The chemiluminescent signal were detected using Chemi-Lumi One Super or Ultra (Nacalai Tesque).

### 2.5 EpiPlex RNA assay

Epitranscriptomic profiling was conducted using AlidaBio’s EpiPlex RNA Mod Encoding kit (P/N 100108) using 2.5 µg purified RNA as input. The RNA was treated with the included DNase I according to protocol directions. Prior to mod enrichment onto EpiPlex beads, the purified RNA was mixed with spike-in controls, fragmented, end-repaired and ligated to an Illumina P7 adapter. Approximately 90% of the prepared RNA fragments were captured onto bead surfaces by mod-specific RNA binders. The remaining 10% were prepared into non-enriched solution control libraries by analogous chemistry. At the bead surfaces, captured RNA fragments are randomly primed by proximally anchored adapters containing Illumina P5 adapters, a unique molecular identifier (UMI) and an MBC (modification barcode) which was extended into cDNA by a reverse transcriptase. Sample indices were added to each sample during PCR using EpiPlex Unique Dual Index Primer Kit (P/N 224001). Libraries were sized on the Agilent TapeStation and quantified by Qubit. Separate library pools containing equal representation of each library were made for bead enriched and solution control libraries. Ribosomal RNA was depleted from each pool using Biorad SEQuoia RiboDepletion Kit (P/N 17006487) according to manufacturer’s directions. Each library was sequenced to ∼25M reads on an Illumina NextSeq1000 using an X-LEAP SBS 200 cycle kit (P/N 20100986).

### 2.6 Peak Calling

Reads were processed using EpiScout, a proprietary RNA modification detection pipeline developed by AlidaBio. Briefly, raw read quality was initially determined using FastQC. MBC and UMI barcodes were then extracted from read sequences, while low-quality bases were trimmed out.

Trimmed reads with length <30bp were removed. Clean reads were then mapped to the human genome (GRCH38.p14) using STAR (29), as well as to our internal spike-in control references using Bowtie2 (30). Genome mapped reads were split by MBC and deduplicated according to MBC and UMI barcodes using samtools and a custom Python script (31).

Peak calling was performed using a custom Python script on a per-modification basis. In brief, peaks were called in a signal-processing manner using the enrichment sample as signal and solution control as a measure of background. Peak regions and fold enrichment were determined using the Python package scipy find_peaks and scaled according to the exogenous spike-in controls’ enrichment factor resulting in final peak fold-enrichment (32). To identify high-confidence peaks, we employed a dynamic filtering technique based on the decay of fold-enrichment in false positive peaks as determined by reproducibility. Peaks which passed this cutoff, along with a minimum depth requirement of 5 reads in the enrichment sample, were considered to be “high-confidence”. After filtering, peaks were annotated using the GRCh38.p14 GTF and fasta files using a custom Python script. Differential analysis was performed using bedtools and a custom Python script. Gene clustering of peaks was performed using the Python package sklearn AgglomerativeClustering (33). Aggregated z-scores were then clustered using Euclidean affinity and ‘ward’ linkage algorithms.

### 2.7 Variant Calling

To accurately identify A-to-G variants captured within EpiPlex inosine peak calls, we used the BCFtools commands mpileup, norm, call, and filter with the mpileup parameters (--full-BAQ, --max-depth 20000), call parameters (-mA), and filter criteria (min(FMT/DP)>5)) (33). Only sites which overlapped inosine peak calls and had a variant rate ≥10% were reported.

### 2.8 Motif analysis

Motif discovery using MEME was performed to identify enriched sequence motifs within inosine peak regions detected in DNA repair genes. The default parameters were used, except that the number of motifs to find was set to 10. To assess whether the identified motifs correspond to known repetitive elements, homology searches were conducted against the Dfam database using *Homo sapiens* as the reference organism.

### 2.9 RNA sequencing and data analysis of gene expression profiles

Total RNA was extracted from 2×10^6^ cells using using the NucleoSpin RNA Plus Kit (Macherey-Nagel, Düren, Germany) following the manufacturer’s instructions. RNA-seq libraries were prepared using the TruSeq Stranded mRNA LT Sample Prep Kit (Illumina, San Diego, CA) according to the manufacturer’s protocol. Sequencing was conducted on the NovaSeq X Plus in 100 bp Paired End mode, using the NovaSeq X series 10B reagent Kit. The raw RNA-seq data underwent adapter trimming and low-quality region removal with Trimmomatic 0.38. The resulting trimmed reads were aligned to the reference genome (GRCh38/hg38) using HISAT2, with transcript assembly conducted using StringTie. Differential expression analysis was carried out with edgeR (v.4.2.2), using raw gene-level read counts. Genes with low expression were excluded by filtering for CPM≧10 across the libraries. After filtering, normalization was conducted using the trimmed mean of M values (TMM) method. Differentially expressed genes (DEGs) were identified using glmQLFTest. Genes with p-value <0.05 and log2 fold change >1 were considered significantly DEGs. Gene ontology analysis of up-regulated and down-regulated DEGs was performed using DAVID.

### 2.10 Gene Ontology and pathway enrichment analysis

Gene ontology enrichment analysis was performed using DAVID, focusing on the Biological Process (GO:BP) category. *Homo sapiens* was selected as the organism, identifiers were provided as OFFICIAL_GENE_SYMBOL. Enriched GO terms with a p-value < 0.05 were considered statistically significant.

## 3 Results

### 3.1 Global profiling of A-to-I RNA editing in human TK6 cells

To identify inosine-modified transcripts in human lymphoblastoid TK6 cells, two independent EpiPlex assays were performed using wild-type cells. The assays detected 2,308 and 2,251 inosine peaks in WT#1 and WT#2, respectively (Table 1 and Table S2). A total of 869 genes were commonly identified in both datasets, demonstrating consistent detection of inosine modifications across biological replicates. Among these 869 genes, A-to-I–modifications were distributed across exons (3 genes), introns (531 genes), 5′-untranslated regions (5′UTRs; 29 genes), 3′UTRs (261 genes), and long non-coding RNAs (lncRNAs; 76 genes) (Fig. 1A). These figures include transcripts with modifications detected in multiple regions.

**Table 1.** Number of inosine- and m^6^A-peaks detected in human TK6 cells by EpiPlex assay.

| Cell line | Number of peaks containing inosine | Number of peaks containing m<sup>6</sup>A |
| --- | --- | --- |
| WT#1 | 2308 | 11040 |
| WT#2 | 2251 | 10786 |
| p150 KO#1 | 736 | 11827 |
| p150 KO#2 | 479 | 9771 |
| p150/p110 KO#1 | 4 | 10884 |
| p150/p110 KO#2 | 2 | 11282 |

**Fig. 1.**
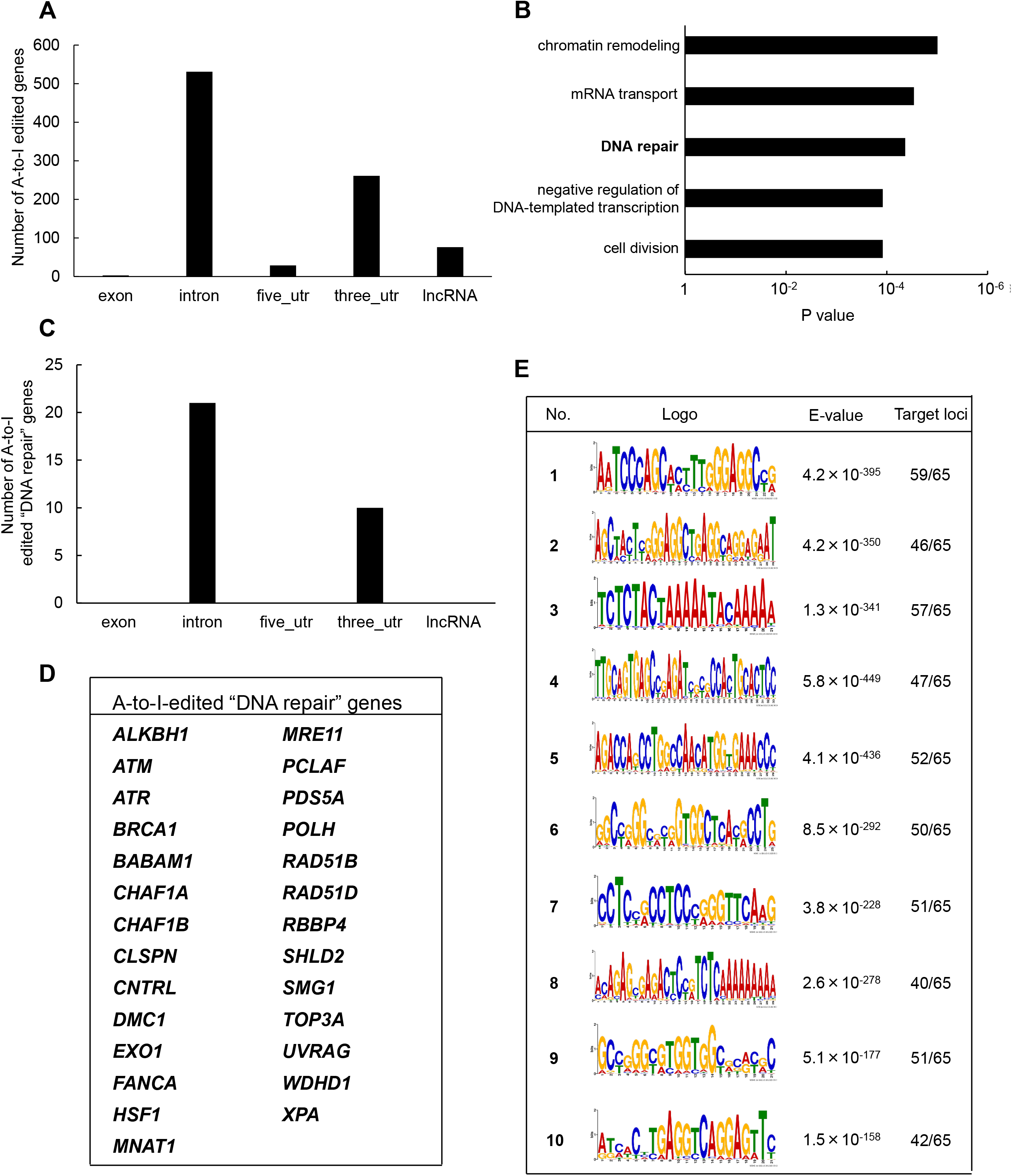
Identification of A-to-I RNA editing sites and associated biological pathways in human TK6 cells. (A) Distribution of A-to-I editing peaks in transcripts expressed in wild-type TK6 cells. Peaks commonly detected across two independent biological replicates are shown. (B) GO enrichment analysis of genes encoding A-to-I-edited RNAs. Statistical significance was determined at p < 0.05, and the top five enriched biological processes are shown. (C) Distribution of A-to-I editing peaks within transcripts of genes involved in DNA repair. (D) List of genes classified under the DNA repair category based on GO analysis. (E) Top ten enriched sequence motifs in A-to-I– edited transcripts of DNA repair-associated genes (MEME analysis). E-values indicate statistical significance. Target loci represent 65 inosine peaks from 27 DNA repair genes.

To explore the biological role of the genes containing A-to-I-modified RNAs, GO enrichment analysis was conducted using the DAVID bioinformatics tool. The analysis revealed significant enrichment of genes involved in “chromatin remodeling,” followed by “mRNA transport,” “DNA repair,” “negative regulation of DNA-templated transcription,” and “cell division” pathways (Fig. 1B). The genes associated with each pathway are listed in Supplementary Table S3. Based on the GO results, 27 genes were classified under DNA repair-related pathways (Fig. 1C). These included factors involved in DNA double-strand break (DSB) repair (i.e., *ATM, ATR, BRCA1, MRE11, TOP3A, EXO1, RAD51B, RAD51D, SHLD2*), nucleotide excision repair (i.e., *XPA*), mismatch repair (i.e., *EXO1*), base excise repair (i.e., *ALKBH1*), interstrand crosslink repair (e.g., *FANCA*), and translesion DNA synthesis (i.e., *POLH*) (Fig. 1D). Among these 27 genes, A-to-I– modifications were located in introns (21 genes) and 3′UTRs (10 genes). To further examine the sequence characteristics of A-to-I–edited sites, motif analysis was performed using MEME (Multiple EM for Motif Elicitation) (34), on 65 inosine peak sequences from the 27 DNA repair-related genes. As shown in Figure 1E, shared sequence motifs were identified across several peak sequences. To determine whether these motifs matched known repetitive elements, homology searches were conducted using the Dfam database. As a result, motif 2 correspond to AluYh7, motif 4 to AluYc, and motif 5 to AluSq4.

### 3.2 Differential contribution of ADAR1 isoforms to A-to-I RNA editing

To assess isoform-specific contributions to inosine modifications, three independent ADAR1 p150-deficient (p150 KO) TK6 cell clones were generated (Fig. 2A and B). These p150 KO cells allowed for distinction between editing mediated by p150 specifically and total ADAR1 activity. Notably, the expression of ADAR1 p110 remained largely unchanged in p150 KO cells (Fig. 2C). To further delineate isoform contributions, ADAR1 p150/p110 deficient (p150/p110 KO) cells were established by disrupting p110 expression in the p150 KO background (Fig. 2B and C). Comprehensively profiling using the EpiPlex assay was conducted on two independent clones each from WT, p150 KO, and p150/p110 KO cells. Inosine peak counts were markedly reduced in the p150 KO#1 and p150 KO#2 clones to 736 and 479, respectively, compared to 2,308 and 2,251 in WT#1 and WT#2 (Table 1 and Supplementary Table S2). In contrast, only 4 and 2 peaks were detected in p150/p110 KO#1 and p150/p110 KO#2, respectively, indicating a 73.4% reduction in p150 KO cells and a 99.9% reduction in p150/p110 relative to WT. Notably, m6A peak counts did not significantly differ across WT and KO lines. To capture a broader landscape of A-to-I editing and determine p150-dependency, clustering analysis was performed on transcripts containing editing sites in at least one WT sample (Supplementary Table S2). As shown in Figure 2D and Table S4, clustering categorized 839 genes as strongly p150-dependent, 303 genes as moderately p150-dependent (reduced in p150 KO), and 176 genes as p150-independent (similar or increased in p150 KO). Additionally, 58 genes exhibited new inosine peaks in p150 KO cells, suggesting a complex regulatory landscape.

**Fig. 2.**
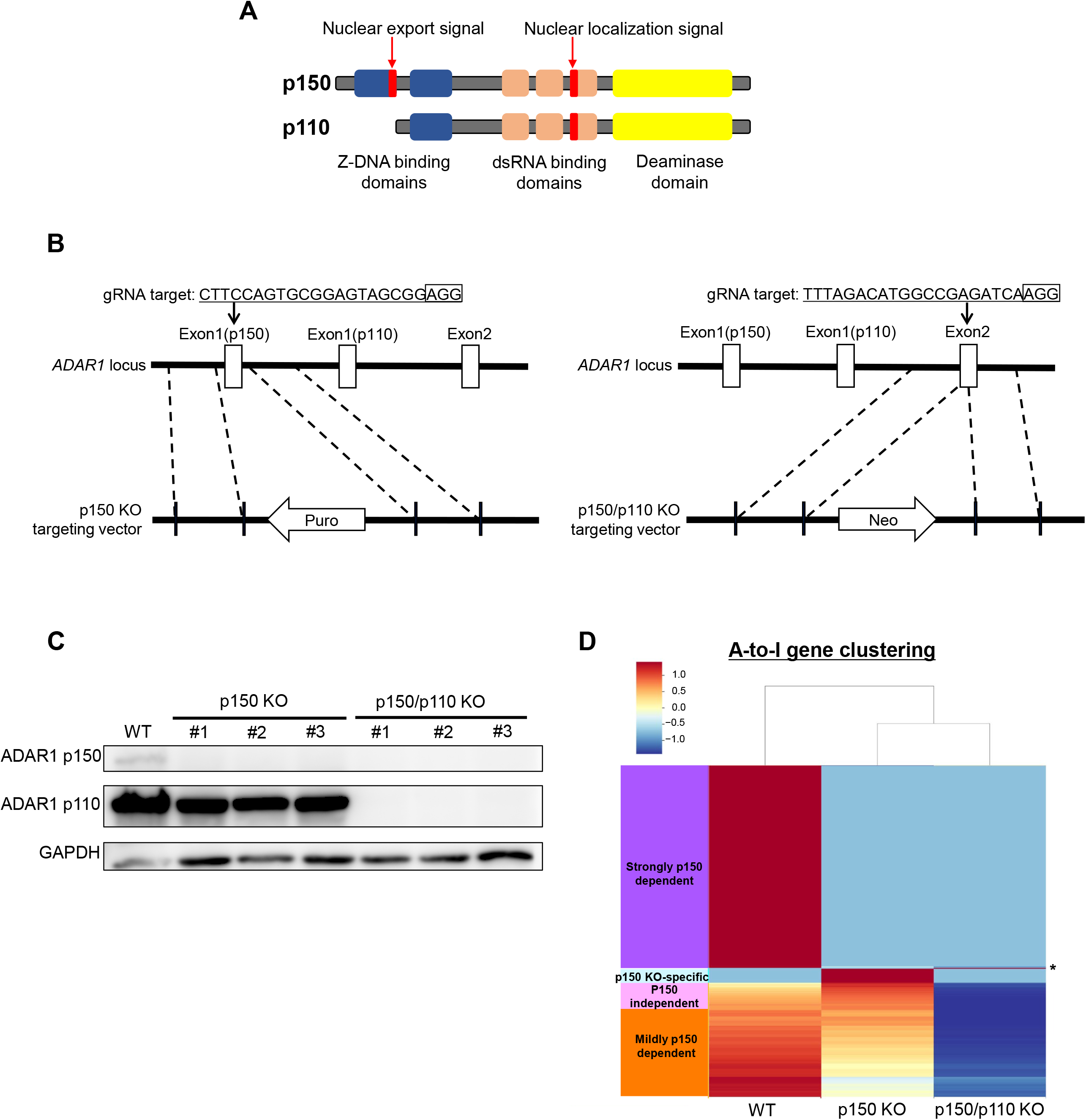
Generation of ADAR1-deficient cell lines for isoform-specific A-to-I RNA editing analysis. (A) Domain structure of the two ADAR1 isoforms, p150 and p110. (B) Schematic representation of the CRISPR/Cas9-mediated disruption of ADAR1 p150 and p150/p110. The target sequence and corresponding vectors containing either a reverse-oriented puromycin- or neomycin- resistance cassette are illustrated. (C) Western blot analysis of ADAR1 isoforms. Whole-cell extracts from WT, p150 KO clones (#1, #2, #3), and p150/p110 KO clones (#1, #2, #3) were resolved by 10% SDS-PAGE. GAPDH was used as a loading control. (D) Clustering analysis of genes exhibiting A- to-I editing. Based on inosine peak scores (fold-enrichment ratio) in WT and p150 KO cells, genes were categorized as strongly p150 dependent (WT/(p150KO+1)≧4; purple), mildly p150 dependent (1<WT/(p150KO+1)<4; orange), p150 independent (WT/(p150KO+1)<1; pink), and newly detected upon p150 loss, i.e., p150KO-specific (WT=0, p150KO>1, cyan). Asterisk (*) indicates the genes categorized as p150/p110 KO-specific (WT=0, p150KO=0, p150/p110KO>1).

Among the 27 DNA repair-related genes with A-to-I-editing, 17 were strongly p150-dependent, 8 were moderately p150-dependent, and 2 were p150-independent (Table 2). To further examine the abundance and distribution of editing sites, transcript visualization using Integrative Genomics Viewer (IGV) was performed. In the 3′UTR, 22 edited bases were found in *ATM* (Fig. 3A and Supplementary Table S5), and 78 in *POLH* (Fig. 3B and Supplementary Table S5). Within introns, 122 edited bases were detected in *POLH* (Fig. 3C and Supplementary Table S5), 22 in *ATR* (Fig. 3D and Supplementary Table S5), 36 in *FANCA* (Fig. 3E and F, and Supplementary Table S5), and 19 in *XPA* (Fig. 3G and Supplementary Table S5). Thus, the extent of editing varied markedly among genes.

**Table 2.** ADAR1 isoform dependency of A-to-I editing in transcripts of DNA repair genes.

| Isoform dependency | Locus of A-to-I editing |  |  |  |  |
| --- | --- | --- | --- | --- | --- |
|  | Intron |  |  | Three_ UTR |  |
| Strongly p150 dependent | ALKBH1 | MNAT1 | UVRAG | ATM | RBBP4 |
|  | BABAM1 | PCLAF | WDHD1 | HSF1 |  |
|  | CHAF1A | RBBP4 | XPA | MRE11 |  |
|  | EXO1 | SHLD2 |  | PCLAF |  |
|  | HSF1 | SMG1 |  | TOP3A |  |
| Mildly p150 dependent | ATR | PDS5A |  | CHAF1B |  |
|  | BRCA1 | POLH |  | CLSPN |  |
|  | DMC1 | RAD51B |  | POLH |  |
| p150 independent | CNTRL |  |  | RAD51D |  |
|  | FANCA |  |  |  |  |

**Fig. 3.**
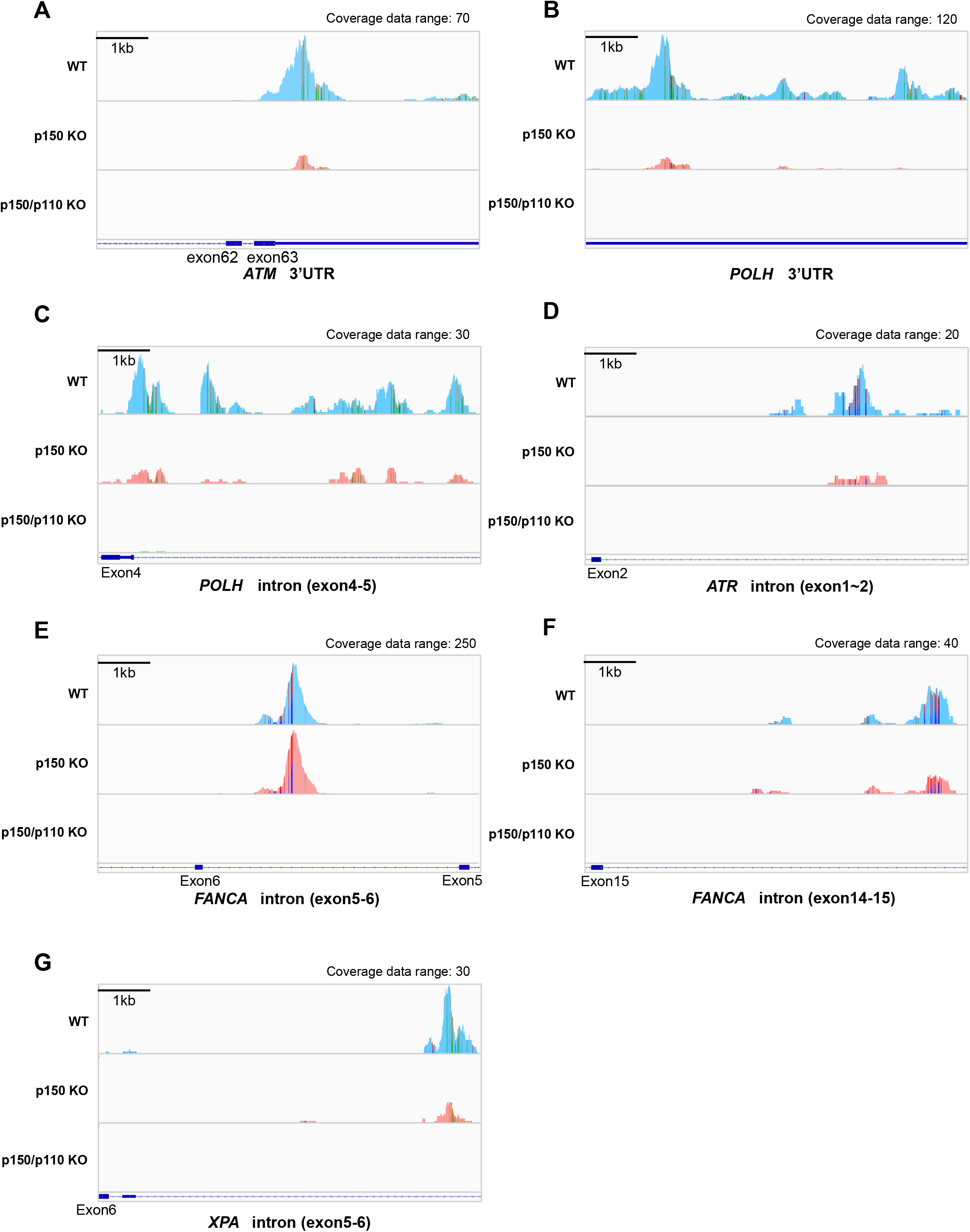
Visualization of A-to-I RNA editing loci in transcripts of DNA repair-related genes in wild-type and ADAR1-deficient TK6 cells. Read coverage profiles are shown for representative DNA repair-associated genes: (A) *ATM* (3′UTR), (B) *POLH* (3′UTR), (C) *POLH* (intron between exon4-5), (D) *ATR* (intron between exon1-2), (E) *FANCA* (intron between exon5-6), (F), *FANCA* (intron between exon14-15), and (G) *XPA* (intron between exon5-6) in wild-type (cyan), p150 KO (pink), and p150/p110 KO cells (green). The y-axis represents read counts and the x-axis indicates genomic coordinates. The scale bar corresponds to 1 kb. “Coverage data range” in the top-right corner of each panel indicates the maximum coverage value. Colored regions within coverage tracks highlight A-to-I RNA editing sites. Strand-specific colors compositions are shown: green (Adenosine) and orange (Guanosine) for forward reads; red (Thymidine) and blue (Cytidine) for reverse reads, facilitating visual estimation of nucleotide ratios at individual position.

### 3.3 ADAR1 deficiency affects *ATM* expression and *XPA* splicing

To evaluate how A-to-I–modifications influence DNA repair gene transcripts, *ATM* (3′UTR modification), *ATR*, and *FANCA* (intronic modifications) were selected as models. mRNA expression levels were measured via qPCR in WT, p150 KO, and p150/p110 KO cells. As shown in Figure 4A, *ATM* expression significantly decreased in p150/p110 KO cells compared to WT, while p150 KO cells showed no significant difference. In contrast, *ATR* and *FANCA* expression levels remained consistent across all three cell lines. Splicing patterns of intron-modified transcripts were also analyzed using IGV analysis of RNA-seq data. Notably, in the *XPA* gene, two novel peaks emerged between exons 5 and 6 in both p150 KO and p150/p110 KO cells compared to WT (Fig. 4B). To validate these splicing events, RT-PCR followed by agarose gel electrophoresis was performed. Additional cDNA bands appeared in the KO cells relative to WT, indicating that ADAR1-mediated A-to-I editing influences *XPA* mRNA splicing (Fig. 4C).

**Fig. 4.**
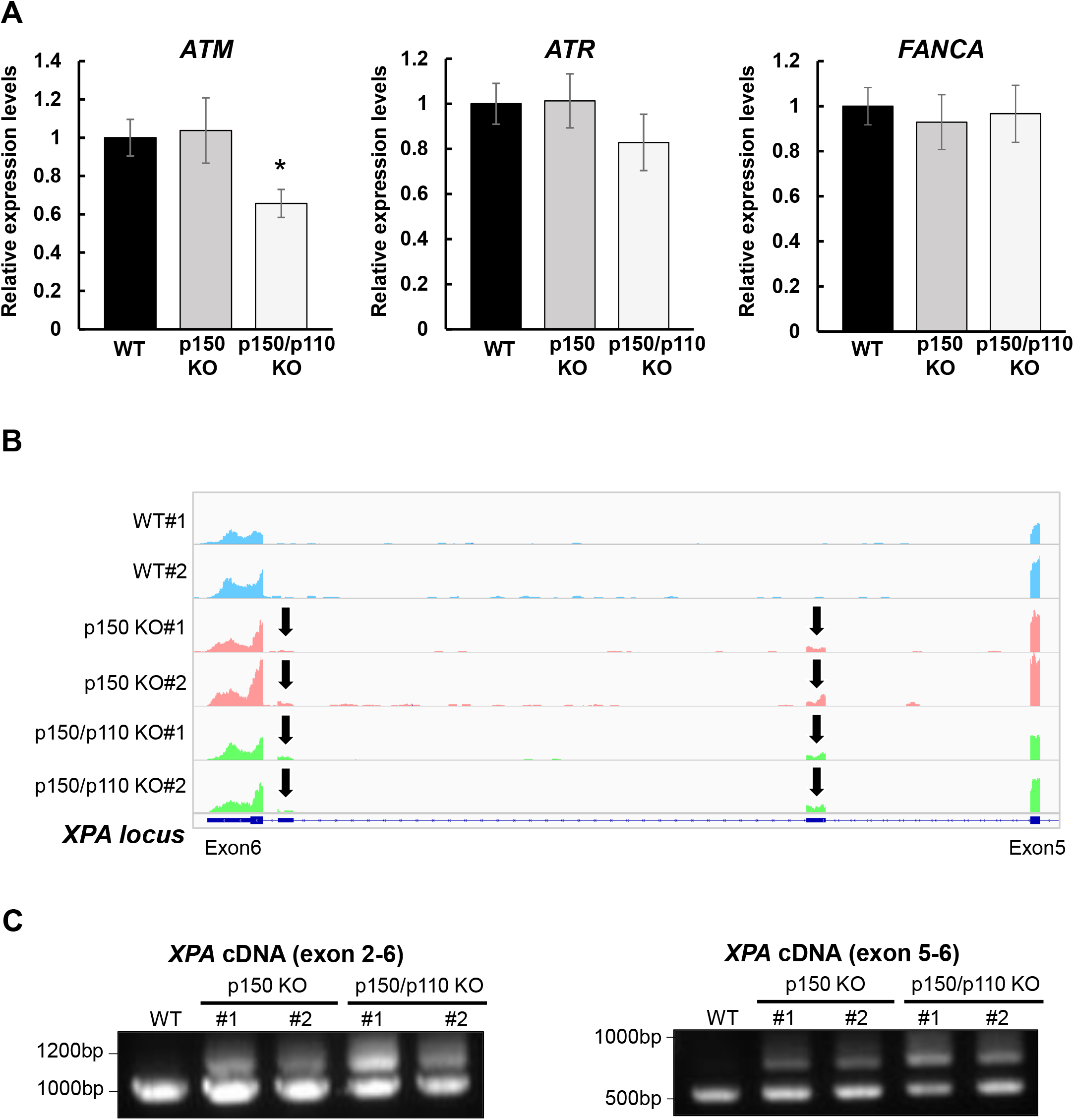
Effects of ADAR1 deficiency on *ATM* expression and *XPA* splicing. (A) Gene expression levels of *ATM, ATR*, and *FANCA* were quantified by RT-qPCR in WT (black), p150 KO (gray), and p150/p110 KO (white) cells. Data represent six independent experiments (n = 6). Statistical significance was determined by Student’s t-test (p < 0.05). (B) RNA-seq read coverage profiles of the *XPA* gene in WT#1, WT#2, p150 KO#1, p150 KO#2, p150/p110 KO#1, and p150/p110 KO#2 cells. Arrows indicate novel splicing peaks between exon 5 and 6 of the main splicing variant. (C) Detection of *XPA* splicing variants. PCR was performed using cDNA from WT#1, p150 KO#1, p150 KO#2, p150/p110 KO#1, and p150/p110 KO#2, with primers targeting exons 2–6 and 5–6. Amplified products were resolved on 1.0% agarose gels. For amplification between exons 2–6, expected band sizes were 946 bp for variant 1 and 1,260 bp for a potential alternative variant. For amplification between exons 5–6, expected sizes were 544 bp for variant 1 and 964 bp for alternative variants.

## 4 Discussion

Recent studies have revealed that A-to-I RNA editing plays a regulatory role in DNA repair mechanisms. For example, in the human breast carcinoma cell line ZR-75-1, A-to-I editing of *ATM* and *POLH* transcripts is enhanced, accompanied by elevated expression levels of these genes, as shown by RNA-seq analysis (24). In human U2OS cells, depletion of ADAR2 has been reported to increase γH2AX levels and the frequency of micronucleus formation, indicating the accumulation of DNA damage (35). Moreover, ADAR1-mediated A-to-I editing influences cellular responses to DNA-damaging agents: knockdown of ADAR1 significantly reduces cell viability following cisplatin treatment in cholangiocarcinoma HuCCT1 and RBE cell lines (25). This phenotype can be rescued by reintroducing wild-type ADAR1, but not by a catalytically inactive mutant (ADAR1 E912A), suggesting that A-to-I RNA editing activity is essential for cisplatin resistance. Thus, A-to-I modification likely plays a critical role in genome stability, and elucidating its broader functions may contribute to novel therapeutic strategies in cancer. Therefore, this study’s comprehensive analysis of the relationship between A-to-I RNA editing and DNA repair pathways provides valuable insight into the physiological role of inosine modifications in genome maintenance.

In this study, the EpiPlex assay, a recently developed for epitranscriptome analysis, was applied to the human lymphoblastoid TK6 cell line, identifying 869 A-to-I-modified transcripts across two independent biological replicates (Supplementary Table S2). The majority of editing sites were located in intronic regions, followed by 3′UTRs (Fig. 1A). This distribution pattern aligns with the global annotation of RNA editing sites annotated in the RADAR database of human transcripts (36, 37). According to GO enrichment analysis, these edited transcripts were significantly enriched in pathways such as “chromatin regulation” and “DNA repair” (Fig. 1B and Supplementary Table S3). Notably, associations between specific biological pathways and RNA modifications appear to be cell type-dependent. For example, in brain tissues, A-to-I editing targets are enriched in transcriptional regulation and neurological disorder–related pathways(38), while in granulosa cells from patients with polycystic ovary syndrome, editing events are associated with the mitotic cell cycle and membrane transport (39). In 14 human cell lines from the ENCODE project, modified RNAs are enriched in cell division, antiviral response, and translational control pathways (40). Collectively, these findings indicate that the functional significance of A-to-I RNA editing varies by tissues and cell type, with a prominent role in DNA repair regulation observed in TK6 cells.

Notably, we identified A-to-I–modifications in mRNAs of 27 DNA repair-related genes, including previously reported targets *ATM* and *POLH* (Fig. 1D)(24). Furthermore, the A-to-I peak sequences of these transcripts shared conserved sequence motifs (Fig. 1E), some of which showed homology to Alu subfamily elements such as AluYh7, AluYc, and AluSq4. This is consistent with earlier reports that A-to-I editing preferentially occurs within Alu elements (41-43), suggesting that A-to-I editing in DNA repair transcripts relies on double-stranded RNA structures formed by Alu-derived sequences. In addition to the DNA repair pathway, GO analysis indicated enrichment in “chromatin remodeling” (Fig. 1B), including genes encoding topoisomerase I (*TOP1*) and histone-modifying enzymes, (*KDM2A, KDM4B, NSD1*, and *NSD2*). These enzymes are known to contribute to the regulation of DNA damage responses (28, 44-46), implying that A-to-I editing may support genome integrity through broader chromatin-based repair mechanisms.

Using the EpiPlex assay to compare wild-type, p150 KO, and p150/p110 KO cell lines, we assessed isoform-specific contributions of ADAR1 to A-to-I editing. Inosine peak levels were reduced by approximately 73.4% in p150 KO cells and by 99.9% in p150/p110 KO cells relative to wild-type cells. In contrast, the number of m6A peaks remained unchanged, suggesting that A-to-I and m6A modifications are independent processes (Fig. 3A). Cluster analysis of A-to-I editing revealed 840 genes as strongly p150-dependent, 302 as moderately p150-dependent, and 176 as p150-independent. Additionally, a distinct subset of genes was edited specifically by the p110 isoform (Fig. 3B). This contrasts with a previous report in HEK293 cells where no p110-specific editing was observed (4), suggesting that ADAR1 editing targets vary by cell type and reflect isoform-specific localization and activity. Consistently, A-to-I editing has been shown to occur in a cell type–specific manner in human brain cells (47). Moreover, under stress conditions, p110 undergoes phosphorylation, promoting transient cytoplasmic translocation (9). Among the DNA repair factors identified in this study, isoform dependency varied across individual genes, implying gene-specific roles for p150 and p110 in A-to-I editing.

Regarding the effects of ADAR1 deficiency on cell viability, our study demonstrated that TK6 cells lacking ADAR1 remain viable (Fig. 2E), in contrast to previous reports where ADAR1 loss induced cell death in HeLa cells. In ALT (alternative lengthening of telomeres)-negative cells like HeLa, ADAR1p110 is required for telomerase-dependent telomere maintenance by editing RNA:DNA hybrids and resolving telomeric R-loops (48): Loss of ADAR1 in these cells destabilizes telomeres, leading to lethal proliferative arrest. In contrast, ALT-positive cells, characterized by heterogeneous telomere lengths, maintain telomere through telomerase-independent mechanisms (49). The viable phenotype of TK6 cells lacking ADAR1 may suggest the presence of alternative telomere maintenance pathways compensating for the loss of inosine-mediated stability. Supporting this idea, a previous study reported that telomerase does not contribute to telomere elongation following ionizing radiation in TK6 cells (50).

Gene expression profiling of ADAR1-deficient cells revealed upregulation of interferon-stimulated genes in both p150 KO and p150/p110 KO cells compared to WT cells (Fig. S1), consistent with studies showing that the loss of A-to-I editing leads to accumulation of endogenous double-stranded RNAs and activation of type I interferon responses (6, 51, 52). RT-qPCR analysis revealed that *ATM* mRNA levels was downregulated in p150/p110 KO cells (Fig. 4A), consistent with previous findings that ADAR1 knockdown breast cancer cells reduces *ATM* expression in ZR-75-1 (24). In contrast, expression levels of *ATR* and *FANCA* remained unchanged (Fig. 4A), indicating gene and locus-specific effects of A-to-I editing on transcript expression. It should be noted that no DNA repair-related genes were detected as DEGs in RNA-seq, indicating that the limited sensitivity of the RNA-seq analysis in quantifying subtle changes in the transcript abundance.

Interestingly, increased expression of novel splicing variants was observed in both p150 KO and p150/p110 KO cells relative to WT (Fig. 4B and C). These alternative splicing events correspond to the pathogenic non-coding transcript variants NR_149093.2 and NR_149094.2 (53). It is plausible that A-to-I editing within central intronic region of *XPA* alters splicing factors binding specificity, thereby reshaping splicing patterns and promoting the production of alternative variants. Supporting this possibility, intronic A-to-I editing has been implicated in splicing regulation at other gene loci. For example, SRSF1 binding to an A-to-I site promotes exonization in the *HNRPLL* gene (54), while SRSF7 binding enhances exon inclusion in *CDDC15* (17). In the *RELL2* gene, ADAR2 binds to a double-stranded RNA structure near the exon, interfering with U2AF65 binding and inducing exon skipping (17). These findings raise the possibility that ADAR1 directly regulates splicing patterns in *XPA* through similar mechanisms.

Previous studies have shown that ADAR1 is crucial for cellular responses to various genotoxic stresses. ADAR1 promotes cell survival following cisplatin treatment through A-to-I editing of the BRCA2 transcript (25). Another recent study found that elevated ADAR1 expression confers chemoresistance to 5-fluorouracil and cisplatin by editing the 3′UTR of the stearoyl-CoA desaturase gene transcript (55). ADAR1 also exhibits anti-apoptotic effects in response to UV-induced stress (9). Under CPT treatment, ADAR1 p110 functions non-catalytically by accumulating at R-loops and recruiting DHX9 and DDX21, thereby enhancing ATR activation (56, 57). Our analysis extends this understanding by demonstrating that ADAR1 broadly edits transcripts encoding genome maintenance factors. These observations suggest that the diverse responses to genotoxic agents under ADAR1 deficiency may result from a combination of impaired A-to-I editing across multiple DNA repair-related transcripts and disruption of ADAR1’s non-catalytic roles, such as R-loop resolution. These findings underscore the multifaceted role of ADAR1 in maintaining genome stability. Further investigations are necessary to clarify the mechanisms by which ADAR1 regulates genome stability, particularly in relation to transcriptomic remodeling and RNA–protein interactions in DNA repair pathways.

## 5 Conflict of Interest

The authors declare that the research was conducted in the absence of any commercial or financial relationships that could be construed as a potential conflict of interest.

## 6 Author Contributions

A.Y., Y.-H.H.-F., Z.J., and A.S. conceptualized the research, A.Y., Y.S., H.T., S.S., Y.-H.H.-F., Z.J., K.U., and A.S. contributed to study discussions and experimental design. A.Y., Y.S., H.T., S.S., Y.-H.H.-F., Z.J., and A.S. performed the experiments and analyzed the data. A.Y and A.S. drafted the manuscript. All authors reviewed and approved the final manuscript.

## 7 Funding

This research was supported by JSPS KAKENHI (22H03748). Additional funding was provided by grants from the Takeda Science Foundation [to A.S. and K.U.].

## 8 Acknowledgments

The authors would like to thank Enago (www.enago.jp) for the English language review.

## 9 Abbreviations

m6A: *N*^6^-methyladenosine
5-mC: 5-methylcytidine
Ψ: pseudouridine
A-to-I: adenosine-to-Inosine
ADAR: Adenosine Deaminase Acting on RNA
dsRNA: double-stranded RNA
3′UTRs: 3′ untranslated regions
sgRNA: single-guide RNA
ALT: Alternative lengthening of telomeres
DEG: Differentially expressed genes
DSB: Double-strand break
GO: Gene Ontology
IGV: Integrative Genomics Viewer
MEME: Multiple EM for Motif Elicitation
UMI: Unique molecular identifier

## 10 Supplementary Material

## 11 Data Availability Statement

The datasets generated for this study have been deposited in the DDBJ with accession number PRJDB35672 (https://www.ncbi.nlm.nih.gov/bioproject/PRJDB35672/).

## Notes

### Competing Interest Statement

The authors have declared no competing interest.

