## Supplementary material for "Epitranscriptome-wide profiling identifies RNA editing events regulated by ADAR1 that are associated with DNA repair mechanisms in human TK6 cells": Figure S1

Fig.S1

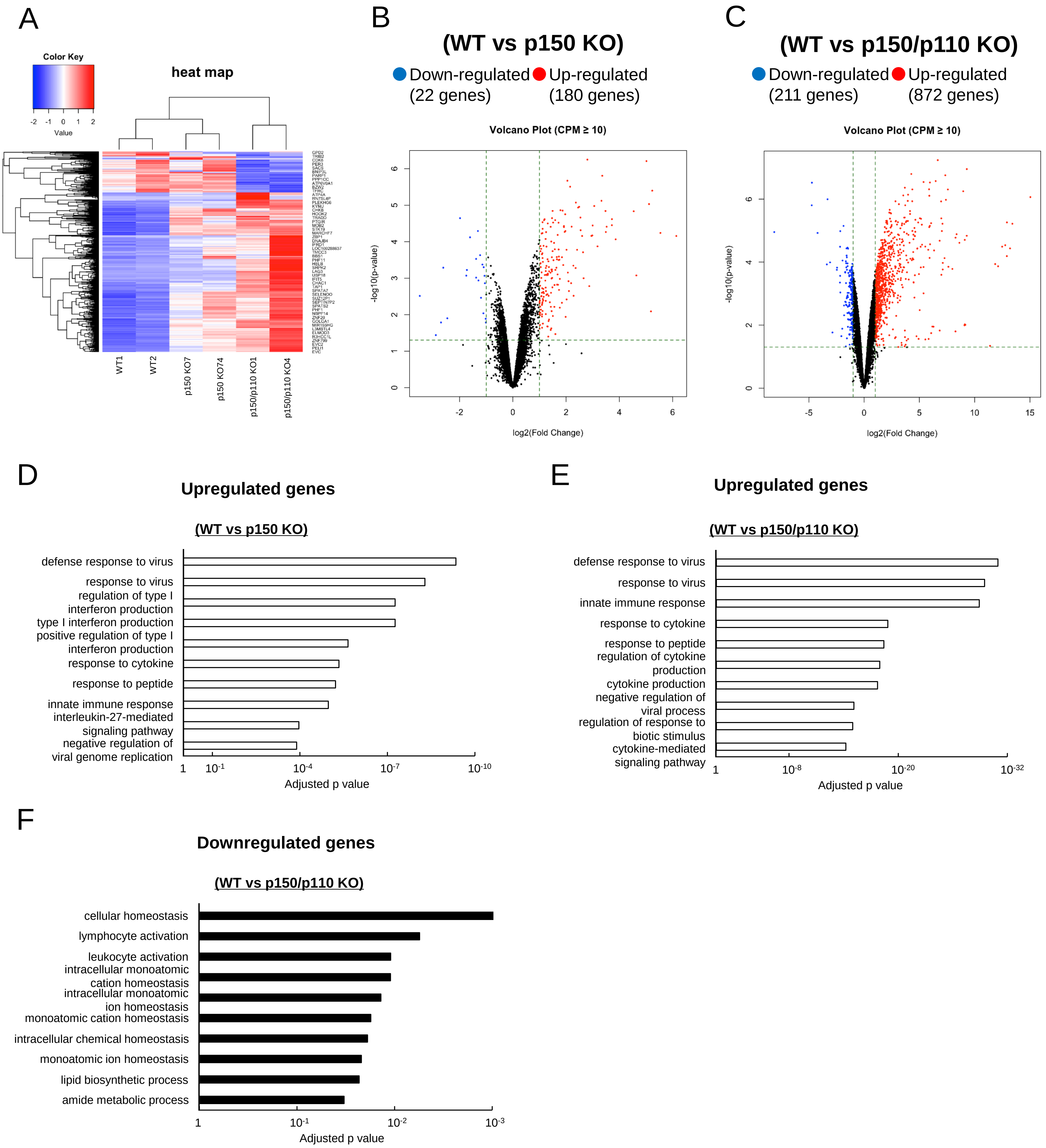


**Figure S1. Differential gene expression between wild-type and ADAR1-deficient TK6 cells.** (A) Heatmap of all differentially expressed genes (DEGs) across WT#1, WT#2, p150 KO#1, p150 KO#2, p150/p110 KO#1, and p150/p110 KO#2 cells. (B) Volcano plot showing DEGs between WT vs p150 KO cells. (C) Volcano plot showing DEGs between WT vs p150/p110 KO cells. (D) GO enrichment analysis of genes upregulated in p150 KO cells compared to WT. (E) GO enrichment analysis of genes upregulated in p150/p110 KO cells compared to WT. (F) GO enrichment analysis of genes downregulated in p150/p110 KO cells compared to WT.
